# Alarm functions of PD-1+ brain resident memory T cells

**DOI:** 10.1101/2024.06.06.597370

**Authors:** Shawn C. Musial, Sierra A. Kleist, Hanna N. Degefu, Myles A. Ford, Tiffany Chen, Jordan F. Isaacs, Vassiliki A. Boussiotis, Alexander G. J. Skorput, Pamela C. Rosato

**Author notes:** Corresponding Author Info.

## Abstract

Resident memory T cells (T_RM_) have been described in barrier tissues as having a ‘sensing and alarm’ function where, upon sensing cognate antigen, they alarm the surrounding tissue and orchestrate local recruitment and activation of immune cells. In the immunologically unique and tightly restricted CNS, it remains unclear if and how brain T_RM_, which express the inhibitory receptor PD-1, alarm the surrounding tissue during antigen re-encounter. Here, we reveal that T_RM_ are sufficient to drive the rapid remodeling of the brain immune landscape through activation of microglia, DCs, NK cells, and B cells, expansion of Tregs, and recruitment of macrophages and monocytic dendritic cells. Moreover, we report that while PD-1 restrains granzyme B expression by reactivated brain T_RM_, it has no effect on cytotoxicity or downstream alarm responses. We conclude that T_RM_ are sufficient to trigger rapid immune activation and recruitment in the CNS and may have an unappreciated role in driving neuroinflammation.

## INTRODUCTION

Tissue resident memory T cells (T_RM_) are distributed throughout the body and are maintained as a self-sustaining and largely non-recirculating population. T_RM_ have been identified in all tissues examined in mice and humans including barrier tissues such as skin, lung, small intestine, and female reproductive tract, and internal organs such as the brain, liver and kidney^1–4^. Foundational studies in barrier tissues demonstrated that T_RM_ provide crucial protection against reinfection by executing a ‘sensing and alarm’ function; they sense reinfection within the tissue and rapidly initiate an alarm response which locally activates and recruits innate and adaptive immunity^3,4^.

Using mouse models of infection and vaccination, studies have shown that T_RM_ seeding in the central nervous system (CNS) is more common than previously thought; brain T_RM_ can be established following local and peripheral infection, and even following peripheral immunization^5–7^. Only recently have CD8+ T_RM_ been identified in human brain, though the antigen specificity of these T cells remains largely unknown^8^. In mice, there is evidence that brain T_RM_ can play both protective and pathogenic roles. Upon pathogen rechallenge, virus-specific brain T_RM_ were shown to be crucial mediators of protection against lethal infection in mice^6,9^. On the other hand, self-antigen specific brain T_RM_ were sufficient to drive neuronal demyelination and tissue destruction^10^. In both conditions, the focus of these studies was on T_RM_-derived cytokine responses and less is known about how T_RM_ might drive immune infiltration and activation in the largely immune restrictive CNS microenvironment.

Given the sensitivity of the brain to inflammation, immune modulation in the CNS likely occurs under multiple mechanisms beyond just restriction of immune migration by the blood-brain barrier. Of note, programmed cell death protein 1 (PD-1) expression has been reported on virus-specific brain T_RM_ across multiple mouse models, and on human brain T_RM_ of unknown antigen specificity^9,11,12^. PD-1 is an inhibitory receptor canonically expressed by activated T cells and further associated with T cell exhaustion in cases of chronic or persisting antigens^13,14^. While much of the initial work regarding brain T_RM_ originated from chronic viral models, more recent work has shown that PD-1 expression is antigen independent and equally characteristic of brain T_RM_ seeded through acute infections^11,12^. To date, PD-1 signaling has only been shown to limit T cell-mediated immunopathology in the brain during the effector phase^11^. As such, it remains unclear how PD-1 signaling might modulate T_RM_ functions in the CNS.

Here, we show that brain T_RM_ can mount a surprisingly fast and robust alarm response, activating microglia, conventional dendritic cells (cDCs), B cells and NK cells, while also recruiting macrophages and monocyte derived dendritic cells (MoDCs) within 48 hours of antigen reencounter. Brain T_RM_ reactivation also triggered expansion of regulatory T cells (Tregs). PD-L1 blockade or genetic deletion of PD-1 resulted in enhanced granzyme B and CD25 expression on brain T_RM_. Despite enhanced reactivation, PD-1 deficient brain T_RM_ were equally capable of cell killing and mounting an alarm response. Together, these data provide a comprehensive analysis of the CNS immune landscape at rest and after T_RM_ reactivation with further insights into the role of PD-1 in regulating brain T_RM_ function. As such, these studies reveal a role for brain T_RM_ in triggering neuroinflammation and may guide immunotherapies targeting unfavorable immune activity in T cell-driven neurological diseases.

## Materials and Methods

### Mice

C57Bl/6J (B6) mice were purchased from The Jackson Laboratory (Bar Harbor, ME) and were maintained in specific-pathogen-free conditions at Dartmouth College. Thy1.1+ OT-I and CD45.1+ OT-I mice were fully backcrossed to C57Bl/6J mice and were maintained in our animal colony. PD-1^−/−^ OT-I mice were generously donated by Dr. Boussiotis at Harvard University^15^. Sample size was chosen on the basis of previous experience. Age and gender matched mice were used for all experiments. Mice used in experiments were 8-20 weeks of age and consisted of both males and females. No sample exclusion criteria was applied. No method of randomization was used during group allocation. All mice were maintained in specific pathogen-free facilities at Dartmouth College under standard housing, husbandry, and dietary conditions according to the Institutional Animal Care and Use Committee (IACUC) and NIH guidelines. All experimental procedures performed were approved by and in accordance with the Institutional Animal Care and Use Committees at Dartmouth College.

### Adoptive transfers and infections

Immune memory mice were generated by transferring 10,000 wild type (WT) Thy1.1+ or CD45.1+ CD8+ OT-I T cells, or CD45.2+ PD-1^−/−^ CD8+ OT-I T cells, from female or male mice into matched naïve 6–8-week-old C57Bl/6 recipients (CD45.2+ or CD45.1+, respectively). One day following transfer, mice were infected intranasally with 5e4 PFU in 30uL of vesicular stomatitis virus expressing chicken ovalbumin (VSV_Ova_)^16^.

### Intravascular staining, lymphocyte isolation, and phenotyping

An intravascular staining method was used to discriminate between cells present in vasculature from cells in the tissue parenchyma as described previously. Mice were injected i.v. with 3ug biotinylated anti-CD8a through tail vein. Three minutes post-injection, animals were sacrificed, and tissues were harvested as described^17^. Mice were euthanized by anesthesia overdose, brains were removed, chopped into small pieces, and incubated with collagenase media (RPMI+5% FBS) containing collagenase type-IV (0.5mg/ml) and DNase (5mg/ml) at 37°C with constant shaking for 30 minutes. After the 30-minute incubation, brain pieces were further dissociated using a GentleMacs dissociator (Miltenyi Biotec) and filtered twice through a 70um mesh. Samples were resuspended in 44% percoll, underlaid with a 67% gradient and spun at 859xG for 20 minutes with the break off. Following percoll spin, the top myelin layer was aspirated, and middle lymphocyte layer was collected into a 15ml conical tube, topped off with RPMI and spun down at 550xG before staining. Isolated lymphocytes were surface and intracellularly stained with indicated antibodies. See Key Resources Table for the complete list of antibodies used for flow cytometry staining. The stained samples were acquired using Cytek Aurora and analyzed with FlowJo software including FlowSom and supervised UMAP plugins^18,19^.

### Intracranial peptide injection, in-vivo antibody treatment

Local T_RM_ reactivation was achieved through intracranial injection of 0.5ug SIINFEKL (Ova) or control (Gp33) peptides in 3ul intracranially as described earlier. Briefly, we made a 0.5-1mm burr hole using a 22-gauge needle without damaging the dura matter. Three microliters of peptide was injected using a Hamilton syringe, into the lateral ventricle (2mm deep, 2mm to the left and 0.5mm south of bregma)^20^. For PD-L1 blockade experiments, 0.2mg of anti-PD-L1 (clone 10F.2H11) antibody was delivered i.v. and 23ug in 3uL intracranially, the next day, an additional 23ug of the blockade was delivered along with 0.5ug Gp33 or Ova peptides in 3uL.

### In-vivo B cell killing assay

B cells were isolated from naïve C57Bl/6 spleens using StemCell EasySep mouse B cell negative selection isolation kit (Cat: 19854A). Isolated B cells were cultured at a concentration of 6e5 cells/1mL with 0.5ug/mL Gp33 or Ova peptide for 20 minutes at 37°C. Cells were washed 3 times to remove excess peptide and resuspended in 500uL of PBS. Ova and Gp33 loaded B cells were stained with CellTrace blue and violet respectively for 5 minutes at 37°C, washed 3 times, combined at a 1:1 ratio and resuspended at a final concentration of 1e6 cells/3uL. Peptide loaded B cells were injected intracranially into mice containing OT-I memory T cells as described earlier.

### Quantification and statistical analysis

All data were analyzed for statistical significance using GraphPad Prism software. Information regarding the tests used, biological replicates, and experimental replicates can be found in the figure legends. Statistical tests included unpaired t tests with Welch correction, one-way ANOVA, two-way ANOVA, and Pearson correlation were performed where appropriate. Bar charts were provided as standard error of mean (± SEM) with bar level representative of the mean variable value. Pearson correlation plot contains fitted linear regression line and R^2^ value. P values < 0.05 were considered significant with asterisks indicated in figure legends.

## RESULTS

### Brain T_RM_ rapidly acquire effector functions following antigen re-encounter

To establish a population of memory T cells, we used a mouse model of acute viral infection with intranasally delivered vesicular stomatitis virus (VSV). VSV is an acute infection that is quickly cleared from mice and leads to establishment of circulating and resident memory T cells throughout the body, including in the brain^5,21^. To enable tracking and manipulation of antigenically defined T cells, we utilized transgenic OT-I CD8+ T cells specific for the SIINFEKL epitope of the model antigen, ovalbumin (Ova). Naïve CD45.1+ or Thy1.1+ OT-I T cells were transferred into adult C57Bl/6 mice intravenously (i.v.) and 24 hours later, intranasally infected with VSV expressing ovalbumin (VSV_Ova_). Mice were rested for a minimum of 30 days to allow for clearance of the virus and establishment of memory T cell populations. These mice will herein be referred to as OT-I memory mice (**Fig. 1A**). To confirm establishment of brain resident memory T cells in this model, as has been previously reported^17^, 30-100 days post-infection we i.v. injected mice with an anti-CD8α antibody 3 minutes prior to euthanasia, which selectively labels T cells in the vasculature but not in tissues^17^. As expected, >90% of T cells in the brain were negative for the i.v. stain and expressed tissue residency markers CD69 and CD103 (**Fig. 1B, C**). Of note, both CD69+/CD103+ and CD69+/CD103-have been shown to be bona fide resident populations in the brain^6,21^. Previous studies have shown that T_RM_ in the brain can execute effector functions such as cytokine secretion following *in vitro* and *in vivo* antigen recall^6,7,22^. We took a reductionist approach to validate this in our model through intracranial (i.c.) injection of cognate Ova peptide. Control mice received an irrelevant peptide (Gp33 epitope from lymphocytic choriomeningitis virus). T_RM_ activation was assessed through direct *ex vivo* staining of effector cytokines and activation molecules 12 and 48 hours later. We found that 12 hours post-peptide injection, brain OT-I upregulated IFNγ, granzyme B and CD25 (IL-2R) (**Fig. 1D-F**). Granzyme B and CD25 upregulation was sustained 48 hours post-peptide (**Fig. 1E, F**), with undetectable levels of IFNγ (data not shown). Moreover, expression of proliferation marker Ki-67 increased on reactivated brain T_RM_ at 48 hours and corresponded to a significant increase in the number of OT-I T cells compared to mice receiving control peptide (**Fig. 1G, H**). These data demonstrate that brain T_RM_ undergo a rapid recall response following local antigen re-encounter with activation sustained over 48 hours.

**Figure 1.**
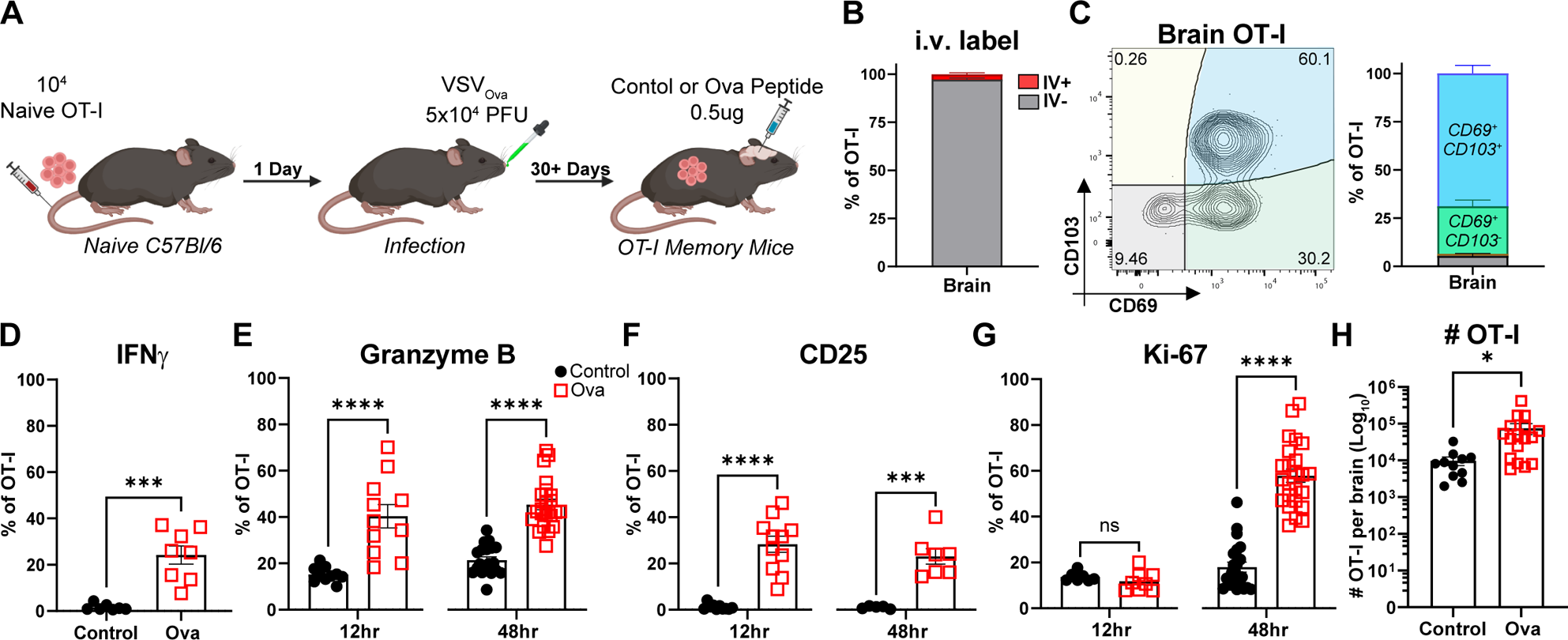
Brain T_RM_ rapidly acquire effector functions following antigen re-encounter. **A)** Schematic of experimental design. Briefly, 10,000 CD45.1+ or Thy1.1+ OT-I T cells were transferred into naïve C57Bl/6 mice followed by intranasal infection with VSV_Ova_. **B)** Bar graph depicting percentage of brain OT-I T cells that are i.v. antibody label stained 30-100 days post infection. **C)** Representative flow plot depicting CD69 and CD103 expression on memory OT-I from the brain (left); Quantified on the right. Gated on i.v.-negative OT-I. **D)** Proportion of IFNg+ OT-I in the brain 12 hours post-peptide. **E-G)** Proportion of granzyme B+ **(E)**, CD25+ **(F)** and Ki-67+ **(G)** OT-I in the brain 12 and 48 hours-post peptide. H) Quantification by flow cytometry of i.v.-negative OT-I T cells in the brain 48 hours post-peptide. **D-H)** Black circles = control peptide; red squares = Ova peptide. Each symbol represents an individual biological replicate pooled from 3-6 independent experiments **(B-H)**. Data are expressed as mean (± SEM) with P values determined by unpaired t-test: ns, not significant, * P< 0.05, ** P < 0.01, *** P < 0.001, **** P < 0.0001.

### PD-1 signaling limits brain T_RM_ reactivation

During the course of our studies, we noticed that brain T_RM_ expressed the inhibitory molecule PD-1, which is canonically expressed on exhausted T cells (**Fig. 2A**). Our observation was consistent with several previous reports noting high PD-1 expression on brain T_RM_ established through multiple models of acute infection^9,11,12^. Given these observations, coupled with our findings regarding effector function following antigen recall (**Fig. 1**), we set out to discern the role for PD-1 signaling on brain T_RM_. Due to the brain’s sensitivity to excessive inflammation and the known role for PD-1 signaling in suppressing T cell responses, we hypothesized PD-1 serves as a regulatory mechanism to modulate the T_RM_ response, thereby limiting T cell-driven neurological damage. To test this, we transferred either wild type (WT) or PD-1^−/−^ OT-I T cells into separate naïve mice followed by intranasal VSV_OVA_ infection. Thirty days post-VSV_Ova_ infection, we assessed OT-I frequencies in the blood and found that genetic deletion of PD-1 resulted in a lower frequency of circulating memory T cells compared to WT (**Fig. 2B**), consistent with previous reports following influenza infection^23^. In contrast, we found that while the frequency of PD-1^−/−^ brain T_RM_ was slightly lower compared to WT, there was no difference in the total number of i.v.-negative OT-I in the brain (**Fig. 2C, D**). Additional analysis of brain T_RM_ phenotype revealed that while nearly all i.v.-negative OT-I expressed CD69, a greater proportion of PD-1^−/−^ OT-I also expressed CD103 (**Fig. 2E**).

**Figure 2.**
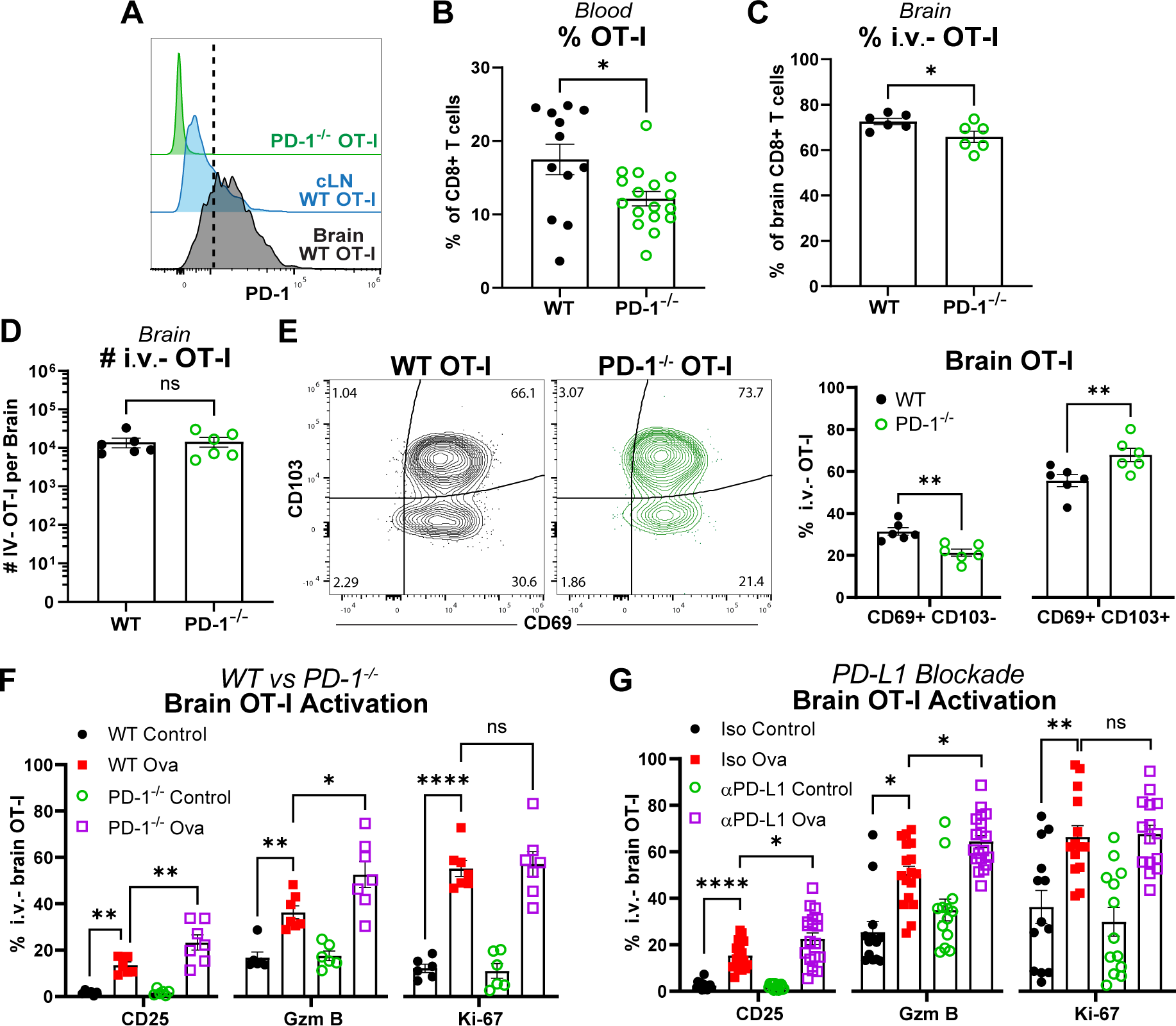
PD-1 signaling limits brain T_RM_ reactivation. **A)** Representative histogram of PD-1 staining across PD-1^−/−^ memory OT-Is compared to WT OT-I T cells from brain and cervical lymph nodes (cLN). **B)** Bar graph depicting percentage of wild type (WT) or PD-1^−/−^ OT-I memory T cells in blood at day 30 post-VSV_Ova_ infection. **C-D)** Proportion of brain OT-I out of total CD8+ T cells **(C)** and absolute number of i.v.-negative OT-I **(D)** from WT or PD-1^−/−^ OT-I recipients. **E)** Representative flow plot depicting CD69 and CD103 expression on WT or PD-1^−/−^ memory OT-I from the brain (left); Quantified on the right. **F)** Proportion of CD25+, granzyme B+, and Ki-67+ WT or PD-1^−/−^ OT-I in the brain 48-hours post-peptide. **G)** Proportion of CD25+, granzyme B+, and Ki-67+ brain OT-I from mice treated with isotype or PD-L1 blockade 48 hours post-peptide. Each symbol represents an individual biological replicate pooled from n = 2 **(B-E)** and n = 4 **(G)** independent experiments. Data are expressed as mean (± SEM) with P values determined by unpaired t-test (**B-E**) or one-way ANOVA (**G**): ns, not significant, * P< 0.05, ** P < 0.01, *** P < 0.001, **** P < 0.0001.

To test the recall response of PD-1^−/−^ brain T_RM_, we injected Ova peptide intracranially, as described in Figure 1. We found that PD-1 deficient brain T_RM_ mounted a greater response as indicated by increases in the frequency of granzyme B and CD25 expressing cells compared to WT (**Fig. 2F**). We saw no difference in proliferation of PD-1^−/−^ brain T_RM_ compared to WT as indicated by Ki-67 staining. As PD-1 signaling has been shown to restrict T cell activation during priming and effector responses to viral infections in other models^23–26^, we repeated our studies using an αPD-L1 blocking antibody after OT-I memory establishment. Blocking PD-L1 resulted in increased expression of granzyme B and CD25, but not Ki-67 in brain T_RM_ compared to isotype IgG (control) recipients, recapitulating our findings with PD-1^−/−^ OT-I T cells (**Fig. 2G**). Taken together, PD-1 signaling is not required for brain T_RM_ establishment, but limits recall responses *in vivo*.

### Brain T_RM_ cytotoxicity is not restricted by PD-1 signaling

The increase in granzyme B expression on PD-1^−/−^ OT-I upon recall led us to hypothesize that PD-1 signaling may restrain T_RM_-driven cytotoxicity in the brain. To assess the direct killing capability of PD-1^−/−^ brain T_RM_ compared to WT we performed an *in vivo* killing assay. Briefly, B cells were isolated from spleens of naïve C57Bl/6 mice and pulsed with either control (Gp33) or reactivating Ova peptide. Peptide loaded cells were labeled using cell trace blue or violet, combined at a 1:1 ratio and injected intracranially into WT or PD-1^−/−^ OT-I memory mice (**Fig. 3A**). Twelve hours later, the frequencies of control and Ova pulsed cells in the brain were assessed by flow cytometry. As expected, we found reduced frequencies of Ova pulsed cells in mice containing WT OT-I brain T_RM_, consistent with published data demonstrating brain T_RM_ cytotoxicity^5^. Interestingly, PD-1^−/−^ brain T_RM_ did not exhibit enhanced killing, as frequencies of Ova pulsed cells were comparable between PD-1^−/−^ and WT OT-I memory mice (**Fig. 3B, C**). Overall, our data suggests that, while PD-1 signaling restricts brain T_RM_ expression of granzyme B and CD25, it does not limit their ability to kill target cells.

**Figure 3.**
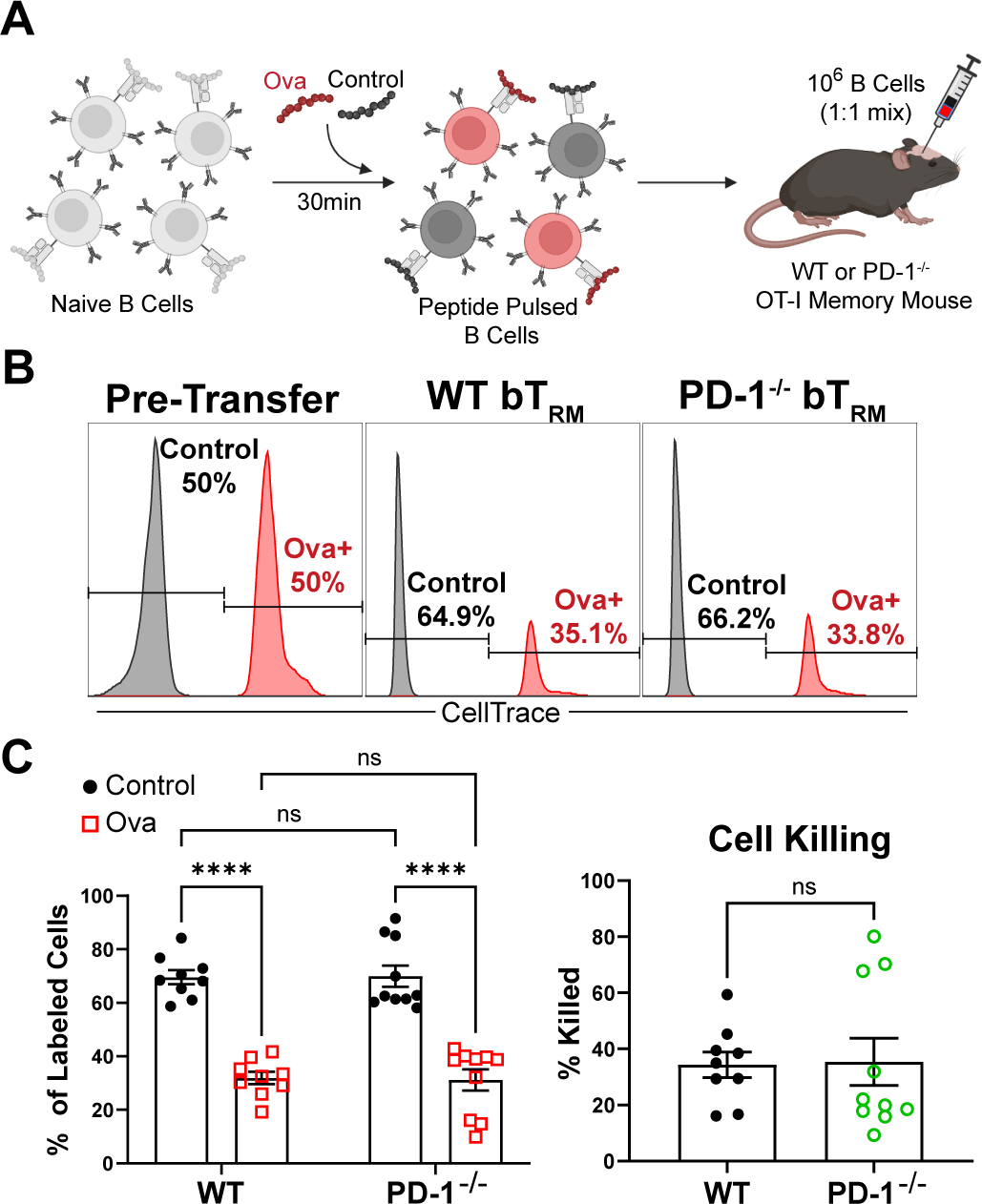
Brain T_RM_ cytotoxicity is not restricted by PD-1 signaling. **A)** Schematic of in-vivo killing assay experimental design. Briefly, naïve B cells were isolated from the spleens of C57Bl/6 mice and incubated with control (Gp33) or Ova (SIINFEKL) peptide, mixed at a 1:1 ratio and injected intracranially (i.c.) into WT or PD-1^−/−^ OT-I memory mice. **B)** Representative histograms of control versus Ova loaded cell ratio pre-transfer and 12-hours later in the brain of WT or PD-1^−/−^ OT-I memory mice. **C)** Bar graph depicting the ratio of control to Ova-loaded cells remaining in the brain after 12 hours (left); Quantification of percentage of killed ova-loaded B cells (right). Each symbol represents an individual biological replicate pooled from n = 2 independent experiments. Data are expressed as mean (± SEM) with P values determined by two-way ANOVA or unpaired t-test: ns, not significant, * P< 0.05, ** P < 0.01, *** P < 0.001, **** P < 0.0001.

### Brain T_RM_ reactivation induces expansion and activation of NK cells, CD4+ T cells, and Tregs

We next used high dimensional spectral cytometry to assess the impact of T_RM_ reactivation on remodeling of CNS immunity 48 hours after control or Ova peptide injection. By reactivating with locally delivered peptide in the absence of infection, we can attribute all downstream immune activation specifically to OT-I T cells in the brain. Using a 27-color panel, we were able to identify 8 discrete immune cell populations in the brain including CD4+ and CD8+ T cells, OT-I T_RM_, NK cells, B cells, dendritic cells (DCs), macrophages, and neutrophils (**Fig 4A. Supplemental Fig. 1)**. Reactivation of OT-I T_RM_ led to broad upregulation of CD69 along with the selective emergence of granzyme B and proliferation marker Ki-67. In addition to increased OT-I T cell numbers, we saw expansion of NK and CD4+ T cells, with a 26- and 9-fold increase, respectively, which correlated with a significant increase in Ki-67 expression **(Fig 4. B-D)**. Further assessment of the NK cell compartment revealed that brain T_RM_ reactivation promoted NK cell activation as defined by upregulation of CD69 (**Fig. 4C**).

**Figure 4.**
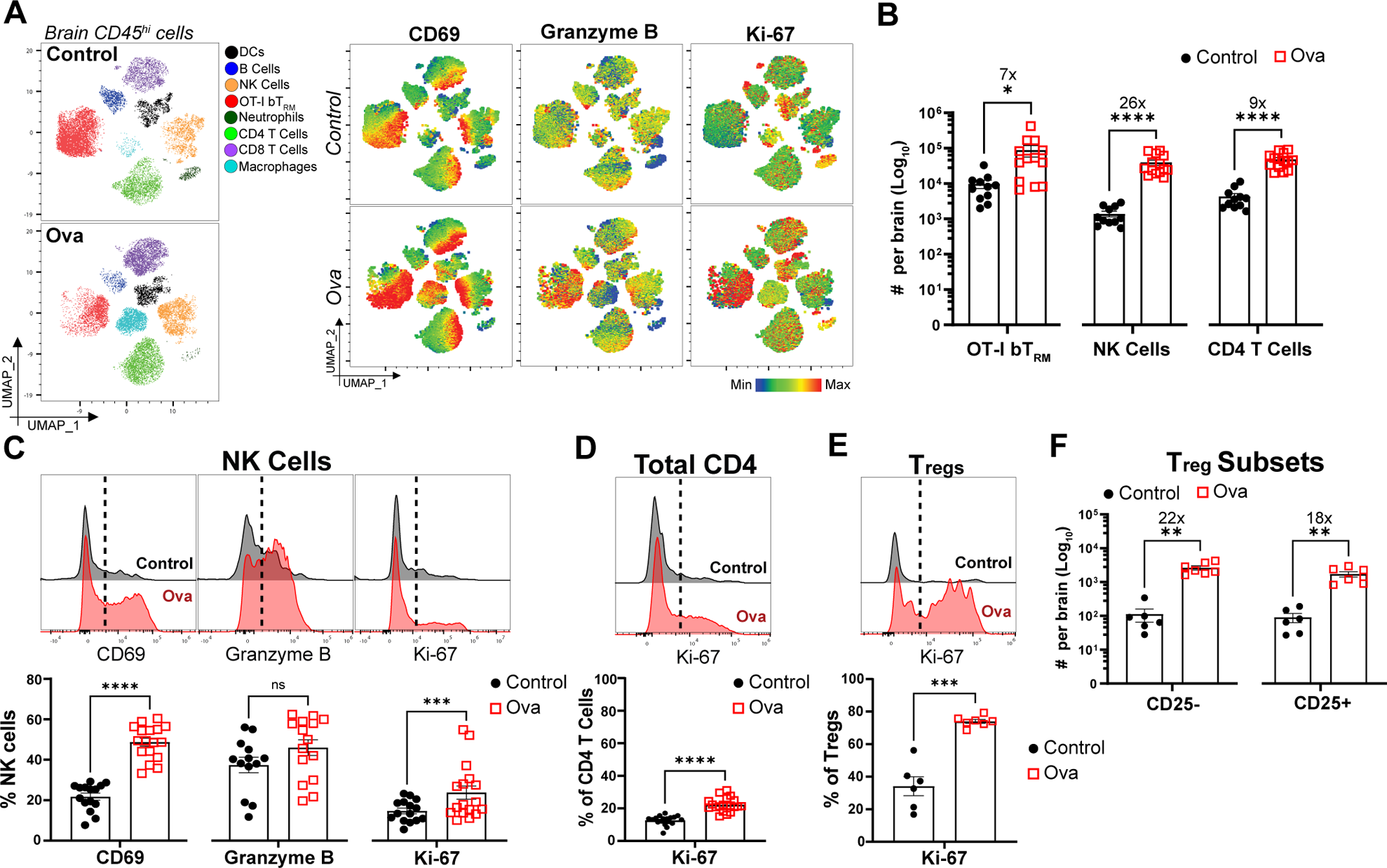
Brain T_RM_ reactivation induces expansion and activation of NK cells, CD4+ T cells, and Tregs. **A)** Supervised UMAP depicting the CD45hi brain immune landscape 48-hours post i.c. control and Ova peptide injections (left); Spectral heatmap overlay illustrating granzyme B, CD69, and Ki-67 expression across the brain immune populations in control and Ova peptide conditions (right). UMAPs are representative and generated with 2 biological replicates downsampled to include 7000 cells each (14000 per condition) and concatenated together. **B)** Absolute numbers of brain OT-I TRM, NK cells, and CD4 T cells 48 hours post control or Ova peptide injections. **C)** Representative histograms of CD69, granzyme B, and Ki-67 expression on brain NK cells 48 hours post control and Ova peptide (top); Quantified below. **D)** Representative histogram of Ki-67 expression on total brain CD4+ T cells 48 hours post i.c. peptide (top); Quantified below. **E)** Representative histogram of brain Foxp3+ Treg Ki-67 expression 48 hours post peptide injection (top); Quantified below. **F)** Absolute counts of CD25- and CD25+ brain Treg subsets 48 hours post peptide. Each symbol represents an individual biological replicate pooled from n = 2-6 independent experiments. Data are expressed as mean (± SEM) with P values determined by unpaired t-test: ns, not significant, * P< 0.05, ** P < 0.01, *** P < 0.001, **** P < 0.0001. See Figure S1 and S2 for supporting data.

CD25 (IL-2R) expression on T cells is often used as a proxy for IL-2 secretion^27^. Our data demonstrating increased CD25 expression on PD-1^−/−^ brain T_RM_ indicates a role for PD-1 in restraining IL-2 production (**Fig. 2**). Given previous reports that T_RM_-derived IL-2 promotes mucosal NK cell activation^28^, we hypothesized that PD-1 restricts T_RM_-induced NK cell activation in the brain. As before, we assessed NK cell activation upon reactivation of PD1^−/−^ OT-I in the brain, or in the presence of αPD-L1 antibody. Contrary to our hypothesis, we found no difference in CD69, granzyme B or Ki-67 expression, or in NK cell numbers when αPD-L1 antibody was delivered at the time of brain T_RM_ reactivation. Genetic deletion of PD-1 from brain OT-I also had no effect on NK cell activation (**Supplemental Fig. 2A-E).**

Our previous analysis was done on total CD4+ T cells and we next wanted to understand if T_RM_ affect Tregs. Foxp3+ Tregs made up approximately 2-8% of total CD4+ T cells in the brain of OT-I memory mice receiving control peptide. Unexpectedly, Tregs demonstrated an increase in proliferation with upwards of 70% of the population positive for Ki-67 staining in response to T_RM_ reactivation (**Fig. 4E**). Consistent with this, we saw a 22- and 18-fold expansion of CD25-negative and CD25+ Foxp3+ Treg subsets, respectively, compared to control (**Fig. 4F**). As before, this was not impacted by genetic deletion of PD-1 on OT-I T cells, though PD-L1 blockade did result in increased Ki-67 expression in total Tregs compared to isotype control (**Supplemental Fig. 2F-I**). In summary, brain T_RM_ trigger expansion of both cytolytic and regulatory immune populations and this is unrestrained by PD-1.

### Brain T_RM_ induce activation, but not expansion, of microglia

We next assessed the effects of brain T_RM_ reactivation on the myeloid compartment, starting with microglia, which are resident macrophages of the brain. Defined here as CD45^mid^/CD11b^+^, microglia did not expand 48 hours following T_RM_ reactivation (**Fig. 5A**). Assessment of microglia from control and Ova recipients by t-distributed stochastic neighbor embedding (tSNE) analysis of flow cytometry data revealed shifts in microglia populations attributed to T_RM_ re-activation (**Fig. 5B**). We found that T_RM_ reactivation resulted in significant upregulation of MHC-II by microglia 48 hours post Ova peptide injection (**Fig. 5C**), characteristic of an activated state^29–31^. Moreover, T_RM_ reactivation led to the emergence of a unique subset of microglia displaying correlated expression of both CD11c and CD86 (**Fig. 5C,D**), both of which have been reported in settings of neuroinflammation and neurological disorders^32–35^. Furthermore, the magnitude of T_RM_-driven microglia activation was not altered under PD-L1 blockade (**Fig. 5E**). Together, these data indicate that brain T_RM_ reactivation drives both microglia activation and emergence of a CD11c and CD86 co-expressing subset.

**Figure 5.**
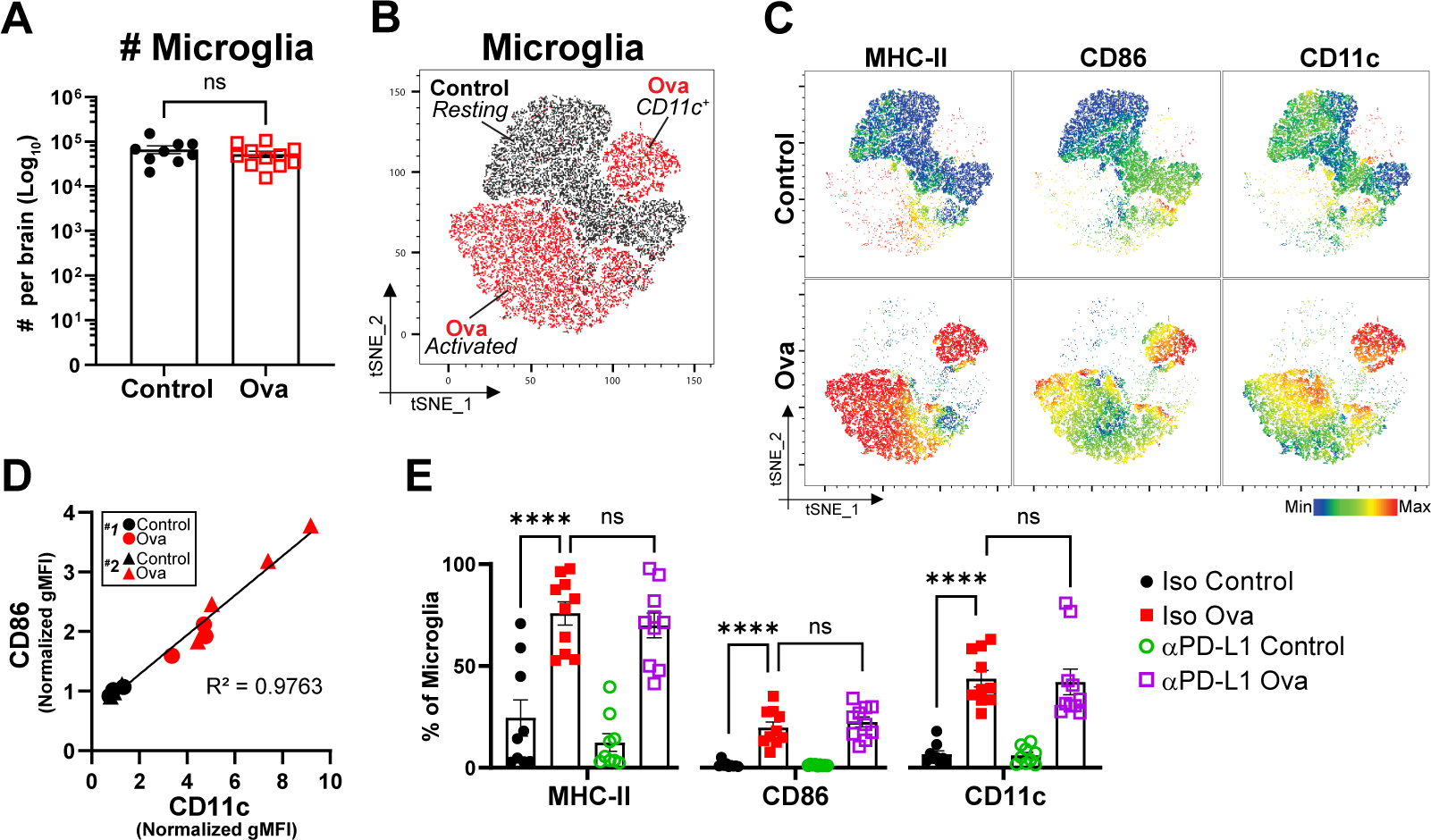
Brain T_RM_ induce activation, but not expansion, of CNS resident microglia. **A)** Absolute numbers of brain microglia 48 hours post control or ova peptide injections. **B)** tSNE plot depicting microglia subsets from control and ova injected brains. tSNE plots are representative and generated with 3 biological replicates downsampled to include 10,000 cells each (30,000 per condition) and concatenated together. **C)** Spectral heatmap overlays of select microglia activation markers across control and ova peptide conditions. **D)** Pearson correlation analysis depicting the relationship between CD11c and CD86 adjusted geometric median fluorescence intensity (gMFI) on microglia from control and ova treated brain. Pooled from n = 2 independent experiments. **E)** Quantification of MHC-II, CD86, and CD11c expression on microglia in recipients treated with either isotype IgG or PD-L1 blockade, 48-hours post i.c. control or ova peptide injection. Each symbol represents an individual biological replicate with n = 2 **(D)** or n= 3 **(A,E)** independent experiments. Data are expressed as mean (± SEM) with P values determined by unpaired t-test (**A**) or one-way ANOVA (**E**): ns, not significant, * P< 0.05, ** P < 0.01, *** P < 0.001, **** P < 0.0001. See Figure S1 for supporting data.

### T_RM_ reactivation induces accumulation of activated B cells in the brain

In addition to microglia, we noted a striking emergence of MHC-II expressing subsets following brain T_RM_ reactivation. As such, we performed unbiased FlowSom clustering on CD45^hi^ MHC-II^+^ cells and identified 6 distinct metaclusters (17 individual clusters) across the brains of both control and Ova peptide treated groups (**Fig. 6A**). Continued breakdown of the CD45^hi^ MHC-II+ brain landscape by experimental group revealed that, while CD19+ B cells are relatively infrequent in the brain, they largely dominated this MHC-II^+^ compartment under control conditions (**Fig. 6B**). Despite their low abundance in the brain, B cells can play a major role in driving disease and protecting again pathogens. Thus, we next examined the impact of brain T_RM_ reactivation on B cell populations. We found a 3-fold increase in the number of B cells in the brain 48-hours following brain T_RM_ reactivation (**Fig. 6C**). Proliferation, as determined by Ki-67 staining, was not elevated (**Fig. 6D**), suggesting that the increased numbers were from infiltrating B cells into the brain and not from expansion of existing populations. We also observed higher frequencies of B cells expressing CD69, CD86 and, interestingly, the macrophage marker, Ly6c (**Fig. 6D**). Moreover, we found that neither αPD-L1 nor genetic deletion of PD-1 on OT-I affected B cell maturation in the brain (**Supplemental Fig. 3**). In summary, T_RM_ reactivation promotes B cell accumulation and activation in the CNS.

**Figure 6.**
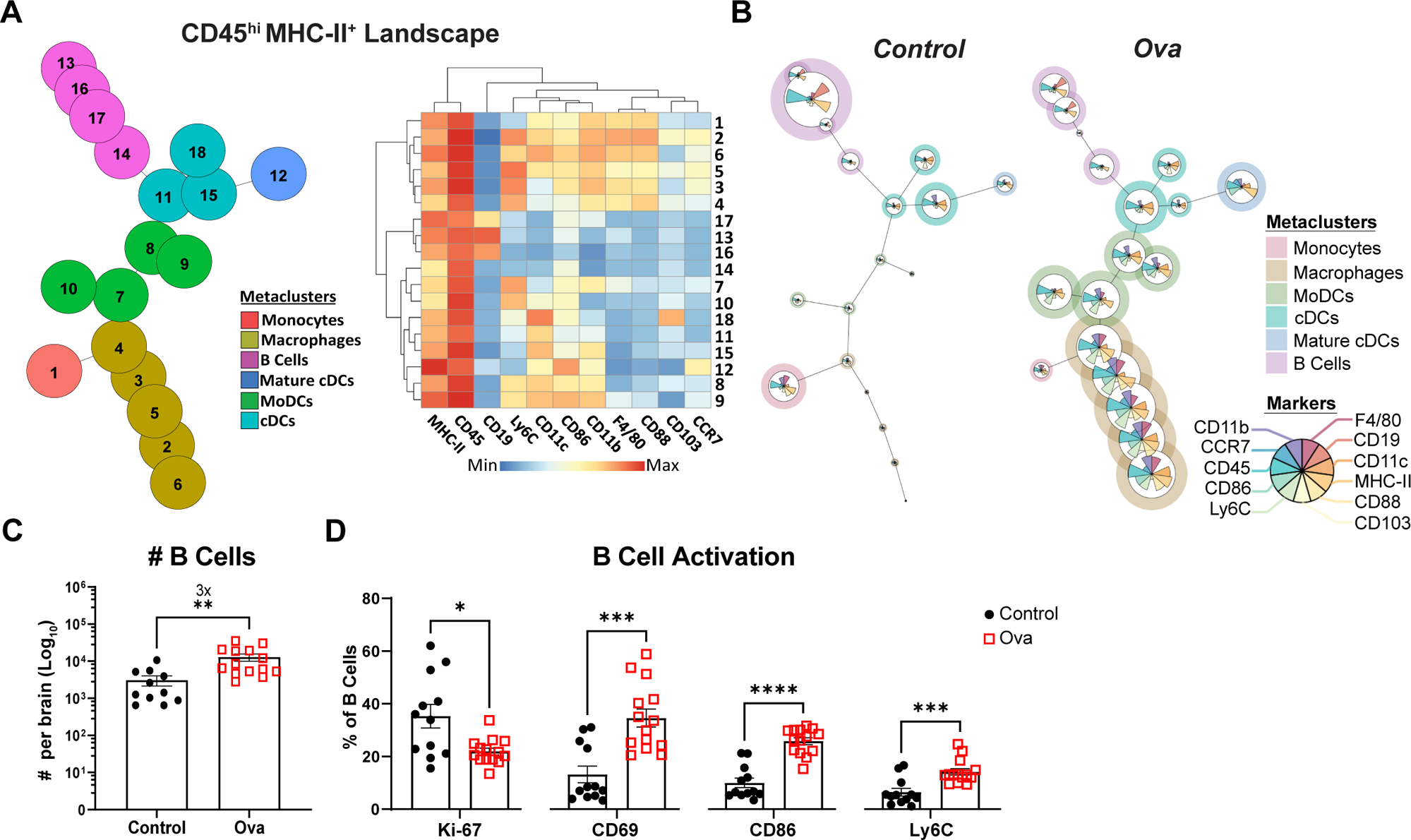
T_RM_ reactivation induces accumulation of activated B cells in the brain. **A)** Concatenated FlowSom dendrogram of CD45hi MHC-II+ immune cells from the brains of both control and Ova recipients, generated utilizing spectral cytometry (left); heatmap of expression pattern of select markers across individual clusters of CD45hi MHC-II+ brain FlowSom analysis (right). A total of 4334 cells were downsampled and concatenated to include 2167 cells per condition. **B)** FlowSom dendrograms of expanded CD45hi MHC-II+ brain subsets in control versus Ova peptide recipients. Dendrograms were generated using the concatenated file containing both control and Ova data, with individual conditions generated using the parent graph. **C)** Absolute numbers of B cells in the brain 48 hours post control or Ova peptide injections. **D)** Frequency of Ki-67+, CD69+, CD86+, and Ly6C+ B cells 48 hours post control and Ova peptide. **C,D)** Each symbol represents an individual biological replicate pooled from n = 4 independent experiments. Data are expressed as mean (± SEM) with P values determined by unpaired t-test: ns, not significant, * P< 0.05, ** P < 0.01, *** P < 0.001, **** P < 0.0001. See Figure S1 and S3 for supporting data

### Brain T_RM_ reactivation induces infiltration and maturation of diverse monocytic cell subsets

T_RM_ reactivation triggered significant remodeling of the CD45^hi^ MHC-II^+^ compartment, characterized by the emergence of monocyte-derived populations (**Fig. 6B**). Notably, we observed infiltrating populations of macrophages and monocyte derived dendritic cells (MoDCs). Macrophage numbers increased 40-fold, constituting roughly 54% of the CD45^hi^ MHC-II^+^ compartment. This was accompanied by a 34-fold increase in MoDCs and 5-fold increase in conventional DCs (cDC) (**Fig. 7A, B**). Interestingly, we report the emergence of mature cDCs upon brain T_RM_ reactivation, as defined by upregulation of both the costimulatory marker CD86 and lymph node homing receptor CCR7 (**Fig. 7A,C**). Finally, we repeated these findings with both PD-1^−/−^ OT-Is and WT OT-Is under PD-L1 blockade and found that the magnitude of the T_RM_-induced myeloid response was unaffected by PD-1 signaling (**Supplemental Fig. 4).** In summary, brain T_RM_ reactivation drives a CNS alarm response that reshapes the brain antigen presenting cell landscape, largely through recruitment of new monocytic populations.

**Figure 7.**
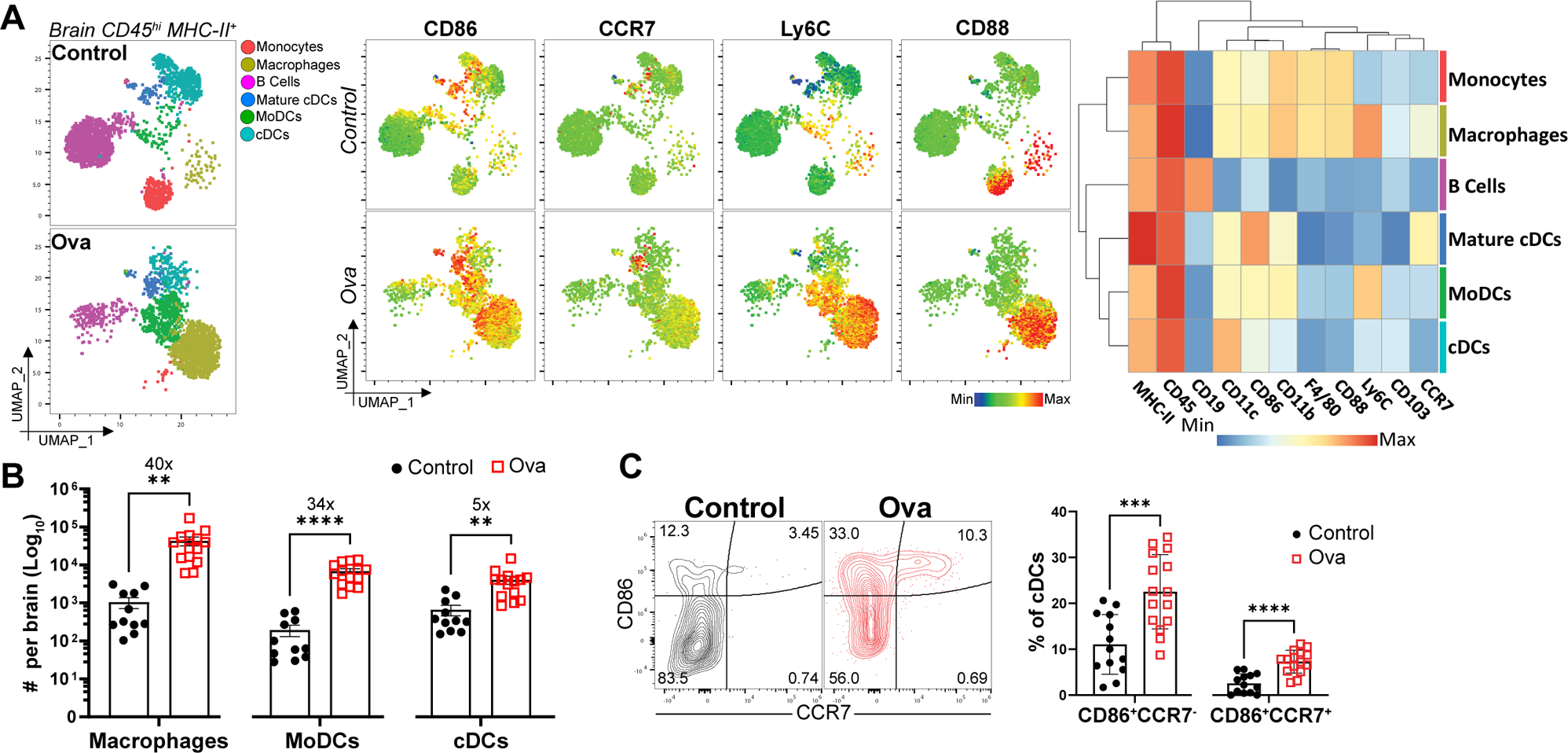
Brain T_RM_ reactivation induces infiltration and maturation of diverse monocytic cell subsets. **A)** Supervised UMAPs of CD45hi MHC-II+ cells from control and Ova injected brains embedded in FlowSom (left); Spectral heatmap overlay of selected surface markers by condition (middle); Heatmap depicting lineage defining markers of FlowSom metaclusters (right). UMAPs are representative and generated with 2 biological replicates downsampled to include 7000 cells each (14000 per condition) and concatenated together. **B)** Absolute number of brain macrophages, MoDCs, cDCs and B cells from control and Ova treated brain 48-hours post i.c. peptide injection, quantified by flow cytometry. **C)** Concatenated flow plots of brain cDC depicting expression of CD86 and CCR7 48-hours post control and ova peptide (left); Quantified to the right. Each symbol represents an individual biological replicate pooled from n = 4 independent experiments. Data are expressed as mean (± SEM) with P values determined by unpaired t-test: ns, not significant, * P< 0.05, ** P < 0.01, *** P < 0.001, **** P < 0.0001. See Figure S1 and S4 for supporting data.

## DISCUSSION

This study defines early T_RM_ recall responses in the brain and demonstrates that T_RM_ reactivation is sufficient to trigger a rapid reshaping of the CNS immune landscape, including activation of local immune populations (NK cells, B cells, microglia, and cDCs), expansion of immune populations (Tregs, CD4+ T cells) and recruitment of new immune subsets (macrophages and MoDCs). Our results place T_RM_ as drivers of neuroinflammation, which has implications for how we contextualize these cells in a disease setting. Several neurologic diseases have been linked to viral infections such as Alzheimer’s disease, epilepsy and recently, multiple sclerosis (MS)^36–38^, however the causative link between infection and disease is unknown. Our studies point to the possibility that reactivation of virus-specific T_RM_ may be an initial trigger for these inflammatory conditions. For example, in the case of MS, we speculate that virus-specific T_RM_ reactivation may exacerbate or trigger disease by enhancing the activation and recruitment of autoreactive B cells in the CNS. Future studies will seek to assess the role T_RM_ play in neurodegeneration and autoimmunity.

The expression of PD-1 on brain T_RM_ across multiple models of viral infection^9,11,12^ might suggest a conserved regulatory mechanism that could modulate T_RM_ reactivation and limit subsequent neurotoxicity driven by the T cell response. While we report enhanced granzyme B and CD25 expression on brain T_RM_ in the absence of PD-1 signaling, this did not translate into enhanced neuroinflammation. Additionally, despite increased granzyme B expression, we found that *in-vivo* killing capacity remained equivalent between WT and PD-1 knockout brain T_RM_. Our findings add to previous work describing a role for PD-1 signaling in restricting salivary gland and lung T_RM_-derived granzyme B, suggesting a conserved regulatory mechanism, rather than a tissue-specific phenomenon^39,40^. Our studies were limited to short-term (within 48hr) assessment of brain T_RM_ reactivation, and it may be that PD-1 signaling plays a role in modulating T_RM_ function to protect against long-term immune mediated damage or cognitive impairment. Moreover, while peptide delivery enabled us to dissect the functional capacity of brain T_RM_, this system lacks the inflammation driven by a viral infection, in addition to the T_RM_-driven inflammation we describe. Thus, it is possible that in the context of viral infection, PD-1 may have a more prominent role in limiting T_RM_-driven immunopathology. It is also possible that brain T_RM_ are regulated by multiple mechanisms, in addition to PD-1, working together and perhaps redundantly to modulate neurotoxicity associated with the recall response; the 20-fold increase in Tregs in the brain following T_RM_ reactivation, for example, may represent a previously unappreciated mechanism by which T_RM_-driven neurotoxicity is kept in check.

The use of PD-1 knockout OT-I T cells in our model raises important considerations regarding the role for PD-1 signaling during the effector phase through memory establishment. As such, a previous study has shown that establishment of memory T cells both in the blood and lung was impaired in the absence of PD-1 signaling following influenza infection. Moreover the few lung T_RM_ that did develop were functionally impaired upon antigen recall^23^. Additionally, Prasad et. al. demonstrated impaired brain T_RM_ establishment following MCMV infection of PD-L1 knockout mice^41^. Our data recapitulates previous reports of reduced circulating memory T cells in the absence of PD-1 signaling, however, it was surprising to find that PD-1^−/−^ brain T_RM_ development was not only unaltered compared to WT T cells, but that PD-1^−/−^ brain T_RM_ mounted a greater recall response. Thus, the differences in T_RM_ developmental requirements for PD-1 signaling may be more nuanced and dependent on the primary infection in addition to tissue of residency.

Despite increases in PD-1^−/−^ brain T_RM_ function, the magnitude of the overall CNS alarm response was equivalent. Our findings build upon previous reports that T cells can activate CNS resident microglia and promote myeloid infiltration^22,42,43^. Brain T_RM_ reactivation prompted the emergence of CD11c+ microglia co-expressing CD86. Although their role in disease remains somewhat contentious, CD11c+ microglia have been reported across neurological disorders like Alzheimer’s and MS^32,44–46^. Our data provides additional insights into the role for brain T_RM_ in initiating the emergence of a specific subset of microglia.

The large increase in macrophage and MoDC populations into the brain 48 hours following T_RM_ reactivation was striking. Though our studies did not deplete circulating cells, it is unlikely that these populations emerged from resident subsets as very few cells could be detected under control conditions. Additionally, both cell types expressed high levels of Ly6C, which is associated with tissue infiltration and inflammation^47,48^. Considering the proposed variance in phagocytic capabilities between microglia and CNS infiltrating macrophages^49–51^, it may be that macrophage infiltration into the CNS following brain T_RM_ reactivation fulfills a specific role in pathogen control, possibly through enhanced phagocytosis. Furthermore, as a previous report observed enhanced antigen presenting capabilities of infiltrating DCs compared to microglia^52^, we hypothesize that MoDCs may serve in the processing and presentation of local antigen for activation of additional T cell responses. One limitation of our study is that all assays involved profiling of the brain as a whole, rather than specific regions such as the meninges or choroid plexus. Future studies will assess immune composition of different brain regions to further understand immune compartmentalization in the CNS driven by T_RM_ reactivation.

Taken together this study demonstrates how brain T_RM_ reactivation drives remodeling of the CNS immune landscape while identifying a role for PD-1 in restraining T_RM_ re-activation. Our results revealed activation of both innate and adaptive immunity in the brain with additional characterization of infiltrating populations post-T_RM_ reactivation. Based on our data, we propose that brain T_RM_ might be repurposed in therapeutic strategies designed to increase immune infiltration in diseases characterized by low immune activity, such as immunologically cold brain tumors^20,53^, or targeted in diseases where T-cell mediated inflammation can be pathogenic, such as multiple sclerosis.

## ACKNOWLEDGEMENTS

We’d like to thank Dr. Vassiliki A. Boussiotis for providing PD-1−/−OT-I mice and David Masopust for providing VSV_Ova_. Additionally, we thank the DartLab Immune Monitoring Core for maintaining cytometers and Dartmouth Animal Care Facility staff members for animal husbandry. Experiment schematics were made using BioRender. This work is supported by NIH grant NIAID K22AI148508 (P.C.R.), R01CA238263 (V.A.B), and Dartmouth Immunology Training Grant T32 AI007363 (S.C.M.).

## AUTHOR CONTRIBUTIONS (CRediT)

Conceptualization, S.C.M. and P.C.R.; Methodology, S.C.M., P.C.R., A.G.J.S., and V.A.B.; Investigation, S.C.M., S.A.K., H.N.D., M.A.F., T.C., and J.F.I.; Resources, S.C.M., M.A.F., V.A.B.; Data curation, S.C.M.; Formal analysis, S.C.M.; Validation, S.C.M.; Visualization, S.C.M. and P.C.R.; Writing – original draft, S.C.M. and P.C.R.; Writing – review & editing, S.C.M, P.C.R., S.A.K., J.F.I., H.N.D., T.C., V.A.B.; Supervision, S.C.M. and P.C.R.; Project administration, P.C.R.; Funding acquisition, P.C.R., V.A.B., S.C.M.

## REFERENCE

1. Schenkel, J. M. & Masopust, D. Tissue-Resident Memory T Cells. Immunity 41, 886–897 (2014).

2. Masopust, D. et al. Dynamic T cell migration program provides resident memory within intestinal epithelium. J. Exp. Med. 207, 553–564 (2010).

3. Schenkel, J. M., Fraser, K. A., Vezys, V. & Masopust, D. Sensing and alarm function of resident memory CD8 + T cells. Nat. Immunol. 14, 509–513 (2013).

4. Wu, T. et al. Lung-resident memory CD8 T cells (TRM) are indispensable for optimal cross-protection against pulmonary virus infection. J. Leukoc. Biol. 95, 215–224 (2014).

5. Wakim, L. M., Woodward-Davis, A. & Bevan, M. J. Memory T cells persisting within the brain after local infection show functional adaptations to their tissue of residence. Proc. Natl. Acad. Sci. 107, 17872–17879 (2010).

6. Urban, S. L. et al. Peripherally induced brain tissue–resident memory CD8 + T cells mediate protection against CNS infection. Nat. Immunol. 21, 938–949 (2020).

7. Steinbach, K. et al. Brain-resident memory T cells represent an autonomous cytotoxic barrier to viral infection. J. Exp. Med. 213, 1571–1587 (2016).

8. Smolders, J. et al. Tissue-resident memory T cells populate the human brain. Nat. Commun. 9, 4593 (2018).

9. Wakim, L. M. et al. The Molecular Signature of Tissue Resident Memory CD8 T Cells Isolated from the Brain. J. Immunol. 189, 3462–3471 (2012).

10. Vincenti, I. et al. Tissue-resident memory CD8+ T cells cooperate with CD4+ T cells to drive compartmentalized immunopathology in the CNS. Sci. Transl. Med. 14, eabl6058 (2022).

11. Schøller, A. S., Nazerai, L., Christensen, J. P. & Thomsen, A. R. Functionally Competent, PD-1+ CD8+ Trm Cells Populate the Brain Following Local Antigen Encounter. Front. Immunol. 11, (2021).

12. Shwetank et al. Maintenance of PD-1 on brain-resident memory CD8 T cells is antigen independent. Immunol. Cell Biol. 95, 953–959 (2017).

13. Freeman, G. J. et al. Engagement of the Pd-1 Immunoinhibitory Receptor by a Novel B7 Family Member Leads to Negative Regulation of Lymphocyte Activation. J. Exp. Med. 192, 1027–1034 (2000).

14. Barber, D. L. et al. Restoring function in exhausted CD8 T cells during chronic viral infection. Nature 439, 682–687 (2006).

15. Strauss, L. et al. Targeted deletion of PD-1 in myeloid cells induces antitumor immunity. Sci. Immunol. 5, (2020).

16. Kim, S. K. et al. Generation of mucosal cytotoxic T cells against soluble protein by tissue-specific environmental and costimulatory signals. Proc. Natl. Acad. Sci. U. S. A. 95, 10814–10819 (1998).

17. Anderson, K. G. et al. Intravascular staining for discrimination of vascular and tissue leukocytes. Nat. Protoc. 9, 209–222 (2014).

18. Van Gassen, S. et al. FlowSOM: Using self-organizing maps for visualization and interpretation of cytometry data. Cytometry A 87, 636–645 (2015).

19. McInnes, L., Healy, J. & Melville, J. UMAP: Uniform Manifold Approximation and Projection for Dimension Reduction. Preprint at 10.48550/arXiv.1802.03426 (2020).

20. Ning, J. et al. Functional virus-specific memory T cells survey glioblastoma. Cancer Immunol. Immunother. 71, 1863–1875 (2022).

21. Nelson, C. E. et al. Robust Iterative Stimulation with Self-Antigens Overcomes CD8+ T Cell Tolerance to Self- and Tumor Antigens. Cell Rep. 28, 3092–3104.e5 (2019).

22. Ayasoufi, K., et al. Brain resident memory T cells rapidly expand and initiate neuroinflammatory responses following CNS viral infection. Brain. Behav. Immun. 112, 51–76 (2023).

23. Pauken, K. E. et al. The PD-1 Pathway Regulates Development and Function of Memory CD8+ T Cells following Respiratory Viral Infection. Cell Rep. 31, 107827 (2020).

24. Lafon, M. et al. Detrimental Contribution of the Immuno-Inhibitor B7-H1 to Rabies Virus Encephalitis1. J. Immunol. 180, 7506–7515 (2008).

25. Erickson, J. J. et al. Viral acute lower respiratory infections impair CD8^+^ T cells through PD-1. J. Clin. Invest. 122, 2967–2982 (2012).

26. Ahn, E., et al. Role of PD-1 during effector CD8 T cell differentiation. Proc. Natl. Acad. Sci. 115, 4749–4754 (2018).

27. Boyman, O. & Sprent, J. The role of interleukin-2 during homeostasis and activation of the immune system. Nat. Rev. Immunol. 12, 180–190 (2012).

28. Schenkel, J. M. et al. Resident memory CD8 T cells trigger protective innate and adaptive immune responses. Science 346, 98–101 (2014).

29. Collawn, J. F. & Benveniste, E. N. Regulation of MHC class II expression in the central nervous system. Microbes Infect. 1, 893–902 (1999).

30. Williams, K., Ulvestad, E. & Antel, J. P. B7/BB-1 antigen expression on adult human microglia studied in vitro and in situ. Eur. J. Immunol. 24, 3031–3037 (1994).

31. Panek, R. B. & Benveniste, E. N. Class II MHC gene expression in microglia. Regulation by the cytokines IFN-gamma, TNF-alpha, and TGF-beta. J. Immunol. 154, 2846–2854 (1995).

32. Wlodarczyk, A., Løbner, M., Cédile, O. & Owens, T. Comparison of microglia and infiltrating CD11c+ cells as antigen presenting cells for T cell proliferation and cytokine response. J. Neuroinflammation 11, 57 (2014).

33. Wlodarczyk, A. et al. Pathologic and Protective Roles for Microglial Subsets and Bone Marrow- and Blood-Derived Myeloid Cells in Central Nervous System Inflammation. Front. Immunol. 0, (2015).

34. Mayrhofer, F. et al. Reduction in CD11c+ microglia correlates with clinical progression in chronic experimental autoimmune demyelination. Neurobiol. Dis. 161, 105556 (2021).

35. Akiyama, H. & McGeer, P. L. Brain microglia constitutively express β-2 integrins. J. Neuroimmunol. 30, 81–93 (1990).

36. Serafini, B., Rosicarelli, B., Veroni, C. & Aloisi, F. Tissue-resident memory T cells in the multiple sclerosis brain and their relationship to Epstein-Barr virus infected B cells. J. Neuroimmunol. 376, 578036 (2023).

37. Bjornevik, K. et al. Longitudinal analysis reveals high prevalence of Epstein-Barr virus associated with multiple sclerosis. Science 375, 296–301 (2022).

38. Wouk, J., Rechenchoski, D. Z., Rodrigues, B. C. D., Ribelato, E. V. & Faccin-Galhardi, L. C. Viral infections and their relationship to neurological disorders. Arch. Virol. 166, 733–753 (2021).

39. Smith, C. J. & Snyder, C. M. Inhibitory Molecules PD-1, CD73 and CD39 Are Expressed by CD8+ T Cells in a Tissue-Dependent Manner and Can Inhibit T Cell Responses to Stimulation. Front. Immunol. 12, (2021).

40. Parks, O. B. et al. PD-1 Impairs CD8+ T Cell Granzyme B Production in Aged Mice during Acute Viral Respiratory Infection. ImmunoHorizons 7, 771–787 (2023).

41. Prasad, S. et al. The PD-1: PD-L1 pathway promotes development of brain-resident memory T cells following acute viral encephalitis. J. Neuroinflammation 14, (2017).

42. Garber, C. et al. T cells promote microglia-mediated synaptic elimination and cognitive dysfunction during recovery from neuropathogenic flaviviruses | Nature Neuroscience. Nat. Neurosci. 22, 1276–1288 (2019).

43. Rosen, S. F. et al. Single-cell RNA transcriptome analysis of CNS immune cells reveals CXCL16/CXCR6 as maintenance factors for tissue-resident T cells that drive synapse elimination. Genome Med. 14, 108 (2022).

44. Benmamar-Badel, A., Owens, T. & Wlodarczyk, A. Protective Microglial Subset in Development, Aging, and Disease: Lessons From Transcriptomic Studies. Front. Immunol. 11, 430 (2020).

45. Holtman, I. R. et al. Induction of a common microglia gene expression signature by aging and neurodegenerative conditions: a co-expression meta-analysis. Acta Neuropathol. Commun. 3, 31 (2015).

46. Remington, L. T., Babcock, A. A., Zehntner, S. P. & Owens, T. Microglial Recruitment, Activation, and Proliferation in Response to Primary Demyelination. Am. J. Pathol. 170, 1713–1724 (2007).

47. Ramachandran, P. et al. Differential Ly-6C expression identifies the recruited macrophage phenotype, which orchestrates the regression of murine liver fibrosis. Proc. Natl. Acad. Sci. U. S. A. 109, E3186–3195 (2012).

48. Li, Y. et al. Occurrences and Functions of Ly6Chi and Ly6Clo Macrophages in Health and Disease. Front. Immunol. 13, 901672 (2022).

49. Yamasaki, R. et al. Differential roles of microglia and monocytes in the inflamed central nervous system. J. Exp. Med. 211, 1533–1549 (2014).

50. Andoh, M. & Koyama, R. Comparative Review of Microglia and Monocytes in CNS Phagocytosis. Cells 10, 2555 (2021).

51. Schilling, M. et al. Predominant phagocytic activity of resident microglia over hematogenous macrophages following transient focal cerebral ischemia: an investigation using green fluorescent protein transgenic bone marrow chimeric mice. Exp. Neurol. 196, 290–297 (2005).

52. Gallizioli, M. et al. Dendritic Cells and Microglia Have Non-redundant Functions in the Inflamed Brain with Protective Effects of Type 1 cDCs. Cell Rep. 33, 108291 (2020).

53. Rosato, P. C. et al. Virus-specific memory T cells populate tumors and can be repurposed for tumor immunotherapy. Nat. Commun. 10, 567 (2019).

