## Supplemental Fig. 1 for "Alarm functions of PD-1+ brain resident memory T cells"

**A**

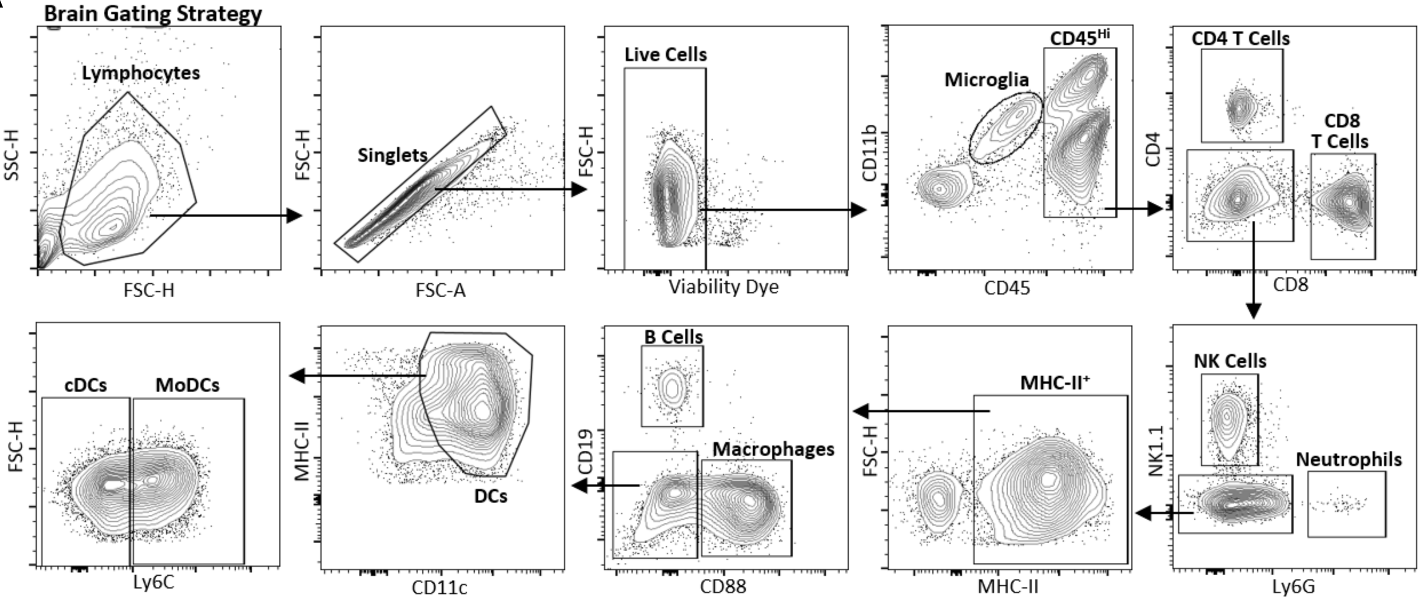


**Supplemental Figure 1. 27-color spectral cytometry brain gating strategy.** Representative gating strategy of immune populations in the brain 48-hours post i.c. ova injection. Immune populations further analyzed in Fig 4 through Fig 7.
