## Supplemental Fig. 2 for "Alarm functions of PD-1+ brain resident memory T cells"

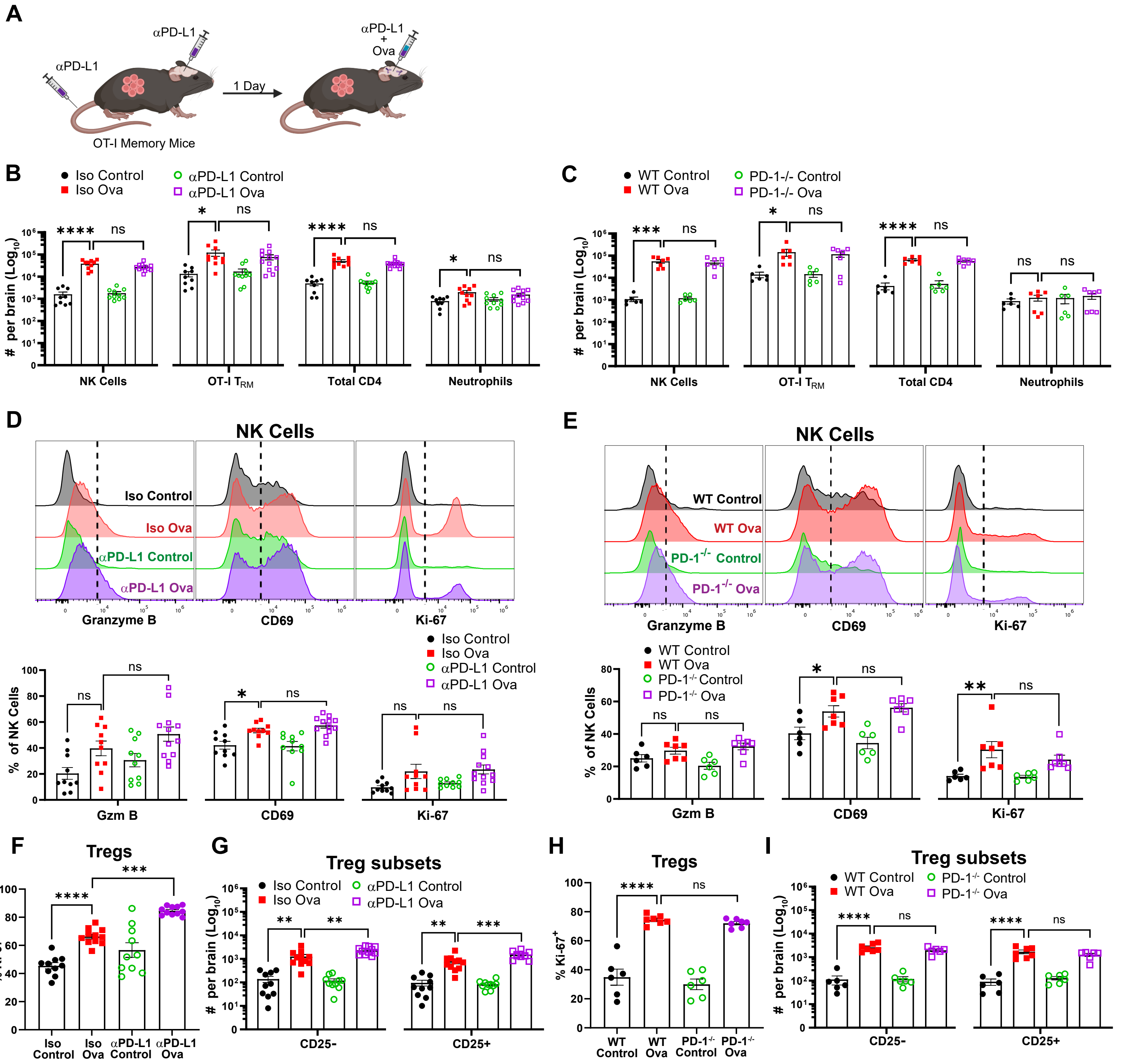


**Supplemental Figure 2. PD-1 signaling does not restrict the magnitude of $T_{RM}$ driven CNS alarm response. A)** Schematic of PD-L1 blockade experimental design. Briefly, OT-I memory mice were established as previously described. 30 days post $VSV_{Ova}$ infection, mice received an i.v. and i.c. injection of either isotype IgG (Iso) or PD-L1 blocking antibody (αPD-L1), followed by an additional i.c. dose on day 31 along with either control or Ova peptide. **B-C)** Number of NK cells, OT-I brain $T_{RM}$, CD4 T cells, and neutrophils in brain of isotype or αPD-L1 recipients **(B)** and PD-1$^{-/-}$ OT-I memory mice **(C)** 48 hours post peptide injection. **D-E)** Representative histograms of granzyme B, CD69, and Ki-67 expression on NK cells from brain of isotype or αPD-L1 treated recipients **(D)** or WT and PD-1$^{-/-}$ OT-I memory mice **(E)** 48 hours post control and ova peptide (top); Graphed below. **F-I)** Frequency of Ki-67+ Tregs **(F),** and number of CD25- and CD25+ Treg subsets from the brain of isotype or αPD-L1 treated recipients **(F,G)** and WT or PD-1$^{-/-}$ OT-I memory mice **(H,I)** 48 hours post peptide. Each symbol represents an individual biological replicate pooled from n = 2 **(C,E,H,I)** and n= 3 **(B,D,F,G)** independent experiments. Data are expressed as mean (± SEM) with P values determined by one-way ANOVA: ns, not significant, * P< 0.05, ** P < 0.01, *** P < 0.001, **** P < 0.0001.
