## Supplemental Fig. 3 for "Alarm functions of PD-1+ brain resident memory T cells"

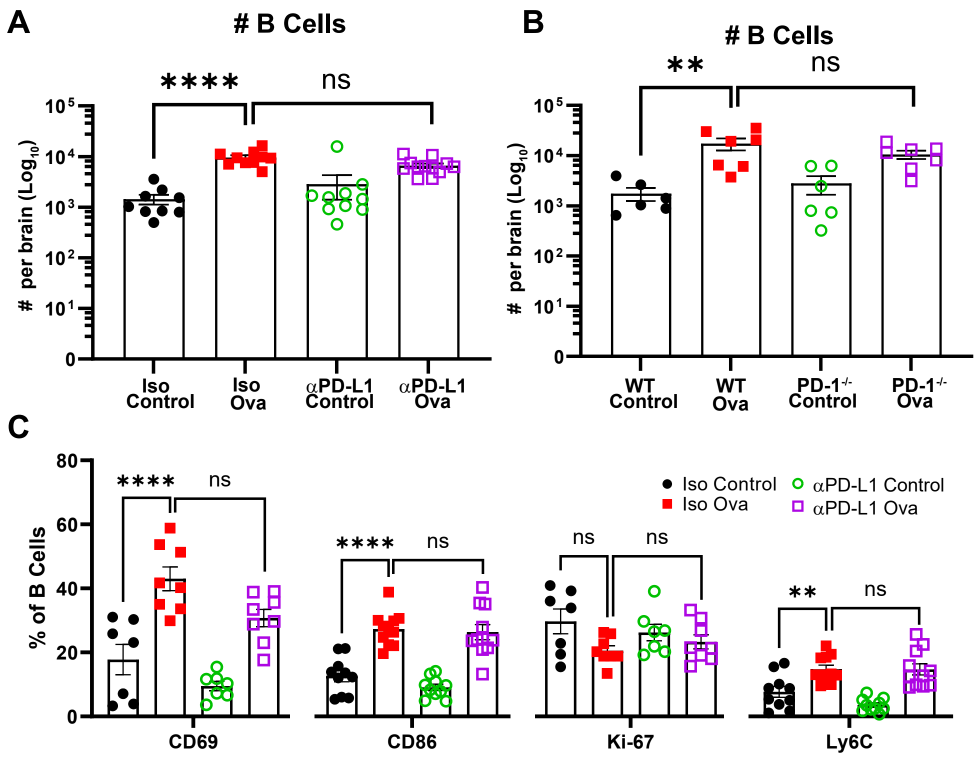


**Supplemental Figure 3. Effects of αPD-L1 treatment and genetic PD-1 deletion on $T_{RM}$-driven B cell maturation in the brain. A-B)** Number of B cells in the brain of isotype or αPD-L1 treated mice **(A)** and WT or PD-1$^{-/-}$ OT-I memory mice **(B)** 48 hours post control or Ova peptide injections. **C)** Frequency of CD69+, CD86+, Ki-67+, and Ly6C+ B cells in the brain of isoptype or αPD-L1 recipients 48 hours post control and ova peptide. Each symbol represents an individual biological replicate pooled from n = 2-3 independent experiments. Data are expressed as mean (± SEM) with P values determined by one-way ANOVA: ns, not significant, * P< 0.05, ** P < 0.01, *** P < 0.001, **** P < 0.0001.
