## Supplemental Fig. 4 for "Alarm functions of PD-1+ brain resident memory T cells"

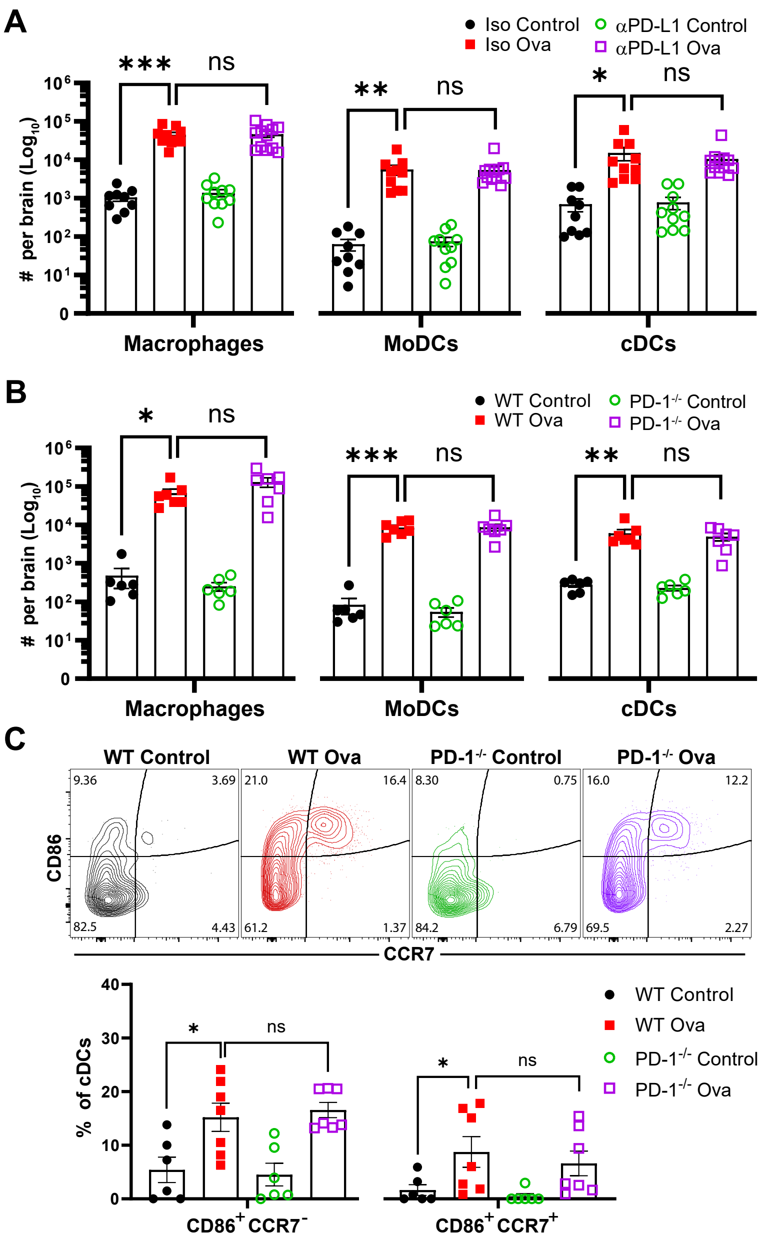


**Supplemental Figure 4. Magnitude of the $T_{RM}$-induced myeloid response is unaffected by PD-1 signaling. A-B)** Number of macrophages, MoDCs, and cDCs in brains of isotype and αPD-L1 treated mice **(A)** or WT or PD-$1^{-/-}$ OT-I memory mice **(B)** 48 hours post peptide injection. **C)** Concatenated flow plot of brain cDC depicting expression of CD86 and CCR7 48-hours post control and ova peptide (top); quantified below. Each symbol represents an individual biological replicate pooled from n = 2-3 independent experiments. Data are expressed as mean (± SEM) with P values determined by one-way ANOVA: ns, not significant, * P< 0.05, ** P < 0.01, *** P < 0.001, **** P < 0.0001.
